# Map segmentation, automated model-building and their application to the Cryo-EM Model Challenge

**DOI:** 10.1101/310268

**Authors:** Thomas C. Terwilliger, Paul D. Adams, Pavel V. Afonine, Oleg V. Sobolev

## Abstract

A recently-developed method for identifying a compact, contiguous region representing the unique part of a density map was applied to 218 cryo-EM maps with resolutions of 4.5 Å or better. The key elements of the segmentation procedure are (1) identification of all regions of density above a threshold and (2) choice of a unique set of these regions, taking symmetry into consideration, that maximize connectivity and compactness. This segmentation approach was then combined with tools for automated map sharpening and model-building to generate models for the 12 maps in the 2016 cryo-EM model challenge in a fully automated manner. The resulting models have completeness from 24% to 82% and RMS distances from reference interpretations of 0.6 Å to 2.1 Å.

## Introduction

In the 2016 Cryo-EM Modeling Challenge (see http://challenges.emdatabank.org/?q=model_challenge; accessed 2017-11-19), a total of 12 maps were supplied to contestants along with reconstruction symmetry and the sequences of the molecules present. One of the goals of the Challenge was to fully interpret such a map given only the map, the symmetry and the sequence information. There are a number of tools being developed by several groups for automated interpretation of cryo-EM maps (DiMaio and Chiu, 2016). These include methods for identification of secondary structure (Jiang et al., 2001; Kong and Ma, 2003; Kong et al., 2004; Baker, Ju and Chiu, 2007), methods for combination of structure-modeling tools such as Rosetta with cryo-EM model-building (Lindert et al., 2012; Wang et al., 2015; Frenz et al., 2017), semi-automated tools for full map interpretation (Baker et al., 2011), and automated tools based on chain-tracing (Chen et al., 2016; Collins and Si, 2017) and template-matching approaches (Zhou, Wang and Wang, 2017).

Prior to the 2016 cryo-EM Model Challenge, we had begun development of software for automatic map sharpening *(phenix.auto_sharpen;* Terwilliger et al., 2018a) and interpretation of density in cryo-EM maps *(phenix.map_to_model;* Terwilliger et al., 2018b) as part of the *Phenix* software package (Adams et al, 2010). It was possible in principle to apply these tools directly to the 2016 Challenge, interpreting an entire map and ignoring the symmetry of the map. It seemed however that it would be more efficient to work with just the unique part of a map. We reasoned that this could be done by identifying a unique part of map that contained a complete molecule, interpreting that part of the map, and then expanding the result using the symmetry in the map to represent the entire map. Powerful tools existed for map segmentation (e.g., Volkmann, 2002; Baker, Chiu and Bajaj, 2006; Yu and Bajaj, 2008; Pintilie et al., 2010), but we wanted to be able to integrate the segmentation and symmetry analysis with automated model-building so that information from model-building could be used to make the final choice of the regions of density representing a single molecular unit. We therefore developed a new *Phenix* tool, *phenix.segment_and_split_map* (Terwilliger et al., 2018b) which could be used for this purpose. Here we describe the application of *phenix.segment_and_split_map* to a set of 218 cryo-EM maps selected to generally represent the unique currently-available cryo-EM maps with resolution of 4.5 Å or better. We then describe map segmentation, sharpening, and model-building (Terwilliger et al., 2018b) applied to the 12 cryo-EM maps in the 2016 Cryo-EM Model Challenge.

## Methods

### Summary of map segmentation

The main goal of our segmentation procedure is to identify the density in a map that corresponds to the unique part of that map. (Note that we use "density" to refer to map values. They can be electron density, electric potential, or any other quantity that is being used to describe the locations of atoms in the map). A secondary goal is to choose this density in such a way that it corresponds as closely as possible to the unique biological unit in the map. Our overall approach to map segmentation is (1) to identify all regions of density above an automatically-determined threshold, and (2) to choose a unique set of density regions that maximizes connectivity and compactness, taking into account the symmetry that is present. By default, the process is repeated with a new threshold after removing the density that has been used in the first iteration. The density threshold for consideration of a region of density is chosen to yield a specific volume fraction (typically 20%) of the region of the macromolecule above the threshold. The map is divided into regions of density above the threshold density, where each region is composed of points above the threshold and that have at least one neighbor above the threshold. A unique set of regions is chosen using the symmetry (if any) supplied by the user and the criteria that the unique set should be as compact and connected as feasible. The details of this segmentation procedure have recently been described (Terwilliger et al., 2018b).

### Symmetry present in a map

We identified symmetry relationships that were applied during map reconstruction using a simplified version of approaches described by Zhang et al., (2012). In many cases the symmetry applied during reconstruction is specified in the EM Data Bank (EMDB, Electron Microscopy Data Bank; Lawson et al., 2016; as for example "I" for icosahedral reconstructions, "C6" for a 6-fold symmetry axis). In others, the symmetry is specified in meta-data associated with the deposited model in the Protein Data Bank (PDB; Bernstein et al., 1977; Berman et al., 2000). In still others, the model deposited in the PDB contains symmetry-related copies which we extracted with the *Phenix* tool *phenix.simple_ncs_from_pdb*. If present, we used symmetry from the deposited models and their meta-data, and if not, we used the information from the EMDB or literature specified in the deposition and the assumption that principal symmetry axes (i.e., screw axes, rotational axes) are generally along the principal axes of the reconstruction to find reconstruction symmetry in the density maps.

### Map-model correlations

We calculated map-model correlations using the *Phenix* tool *phenix.map_model_cc*. This tool identifies the region occupied by the model as all grid points in the target map within a specified distance (typically 3 Å) of an atom in the model. Then it generates a model-based map on the same grid and calculates the correlation of density values between the target map and the model-based map inside the region occupied by the model.

Model-based maps were calculated in reciprocal space using elastic atomic scattering factors of electrons for neutral atoms as described (Colliex et al., 2006, Afonine et al., 2018b). These scattering factors are framed as the sum of Gaussian terms, represent electric potential, and assume that all atoms are independent. These scattering factors do not include the effects of charged residues and therefore they may be substantially incorrect for certain atoms, including phosphates in RNA or DNA and side chains such as aspartate and glutamate. As improved representations of electron scattering expressed as sums of Gaussian terms these become available these can readily be incorporated in the Phenix framework.

### Data used for map segmentation

We selected a group of 218 cryo-EM maps to test our segmentation algorithms. We started with 492 maps we could extract from the EMDB in August of 2017 with simple *Phenix* tools and that were reconstructed at resolutions of 4.5 Å or better. We excluded 91 maps where the resolution in the EMDB and PDB differed by 0.2 Å or more or was not reported, 24 maps where the map-model correlation was less than 0.3, and 16 maps for which the signal-to-noise in map sharpening (Terwilliger et al., 2018a) was less than 3. We then removed map-model pairs that were largely duplications by clustering based on sequence identity using a cutoff of 95% identity and choosing the highest-resolution representative of each group. The sequence identity of two structures was calculated after alignment of each chain in the first structure with the closest-matching chain in the second structure. If either sequence was contained within the other, the identity was considered to be 100%. Otherwise if the lengths of the sequences differed by more than 5%, or the percentage of residues in all chains of the first structure matching a corresponding residue in the second structure was less than 95%, the sequences were considered to be different. Four of the maps in this set were associated with two models, so one map-model pair was set aside for each of these, yielding 218 map-model pairs that were analyzed in this work.

### Evaluating the results of map segmentation by calculation of fraction of molecular unit within the segmented region

We estimated the fraction of the molecular unit within the segmented region of a map from a comparison of map-model correlations. Our method is related to the cross-correlation variation metric described by Zhang et al. (2012) but it is extended to make an estimate of the fraction of the molecular unit that matches the segmented map. The segmented map has values of zero everywhere outside the segmented region of the map. The overall idea is that if the segmented region contains a complete molecular unit, then the map-model correlation between one complete molecular unit and the segmented map will be the same as the map-model correlation with the original map. On the other hand, if the segmented region contains part of one molecular unit and parts of symmetry-related ones, then the map-model correlation between one intact molecular unit and the segmented map will be lower than the correlation to the original map. We use this difference in map correlation to estimate the fraction of a complete molecule that is within the segmented region.

We first calculated the map-model correlation between the original map and a map calculated from single molecular unit extracted from the deposited model of the structure. Then we calculated the map-model correlation between the segmented map and a single molecular unit. The square of the ratio of these correlations is (see below) approximately equal to the fraction of the molecular volume that is within the segmented map. The single molecular unit to compare with the map was chosen to be a set of chains representing the unique part of the deposited model. In cases with symmetry, each symmetry-related choice of molecular unit was considered and the one with the highest map-model correlation was chosen.

The relationship between the map-model correlation for a single molecular unit and the original map compared to the correlation for a molecular unit and a segmented map can be calculated in a straightforward fashion with one assumption. This assumption is that the local map-model correlation for the original map and this single molecular unit is approximately the same everywhere in the region of the model. With this assumption, we can readily calculate the effect of setting all but a fraction *f* of the map density in the region of the model to zero. This corresponds to calculating the map-model correlation of the segmented map to one full molecular unit, where a fraction *f* of the molecular unit is present in the segmented map.

The correlation coefficient CC between two maps with density values represented by D_1_ and D_2_ can be written (after adjusting each map to set the mean density for each to zero so that <D_1_> = <D_2_> =0) as,

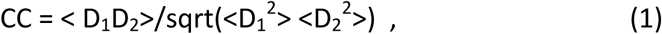

where the calculation in this case is carried out over all the grid points near the model. Now suppose we create a new map D_1_’ in which we set D_1_ to zero at a fraction (1-*f*) of these grid points. Referencing Eq. (1), this means that the values of D_1_’D_2_ and D_1_’^2^ will be zero at all these grid points, but D_2_^2^ will be the same. Assuming then that the values of < D_1_D_2_>, <D_1_^2^>, and <D_2_^2^> are approximately the same everywhere near the model, we can write that the map-model correlation for the segmented map (CC') with all but (1-*f*) of the map set to zero is related to the map-model correlation for the original map (CC) by,

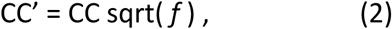

so that *f*, the fraction inside the mask, is approximately given by *f* = CC'^2^/CC^2^.

### Automated model-building

The *Phenix* tool *phenix.map_to_model* has recently been described in detail (Terwilliger et al., 2018c). The inputs required are a map file (CCP4/MRC format, Cheng et al., 2015), a sequence file with the sequences of residues or nucleotides in each unique chain in the structure, and the nominal resolution of the map. If symmetry was used in the reconstruction process, then the symmetry operators can be supplied as well. All other parameters are fully optional and it is normally not necessary for a user to adjust them.

For the model-building described here, the maps, sequence files, symmetry operators, and resolution were all obtained from the 2016 Model Challenge web site at http://challenges.emdatabank.org/?q=model_challenge.

The first step carried out by the map_to_model tool is to automatically sharpen the map with the *phenix.auto_sharpen* tool (Terwilliger et al., 2018a). In this approach the map is sharpened (or blurred) to attempt to simultaneously maximize the level of detail in the map and the connectivity of the map.

The second step is to carry out automatic map segmentation as described above, yielding one map that represents the unique part of the sharpened map along with a set of small maps each representing one small region of connected density (all above a contour level determined automatically during the segmentation process).

The third step is to carry out automatic model-building for each chain type that is represented in the sequence file. This is done for the map representing the unique part of the sharpened map and for each small map. Model-building is done using tools available in *Phenix* that include placement of helices and strands in density of corresponding shapes (Terwilliger, 2010a; Terwilliger, 2010b), tracing density along a chain and replacement with main-chain atoms (Terwilliger, 2010c), placement of short fragments by convolution-based searches followed by extension with 3-residue fragments from structures in the PDB (Terwilliger, 2003; Terwilliger et al., 2018c), and recently-described methods for model-building of RNA that are extensions of these procedures for protein (Terwilliger et al., 2018c).

The fourth step is to combine all the models. The principal method for combining models is to rank all segments (fragments of a model that have no chain breaks) based on map-model correlation, segment length, and secondary structure, then to go through this ranked list and place whatever part of each model does not overlap with a higher-scoring model (Terwilliger et al., 2018c).

After each model is built, after models are combined, and after application of reconstruction symmetry to the final model, each working model is refined with real-space refinement (Afonine et al., 2018a).

### Data used from the Cryo-EM Model Challenge

The maps and reconstruction symmetry used for the 12 cryo-EM maps in the 2016 Cryo-EM Model Challenge were taken from the Model Challenge site at http://challenges.emdatabank.org/?q=model_challenge (accessed 2017-11-19). The Challenge consisted of 8 unique molecules, four of which were associated with two maps at different resolutions, leading to 12 different maps (Table I). Of these maps, most were associated with previously-deposited models that were likely to be more accurate than the ones we built automatically and that were therefore suitable for use as references for the accuracy of our models. For one additional map (groEL, EMDB entry 6422) there was no deposited model, however there is a model for a related structure in the PDB (1ss8) which we offset superimposed on this map and used as a reference. One final structure was recently interpreted (the proteasome structure; Veesler, D., unpublished) and we used that structure as a reference model. We checked the map-model agreement with *phenix.map_model_cc* and these map-model correlations ranged from 0.34 (rather low, supporting only low confidence in the model), to 0.85, suggesting that the model is in good agreement with the map.

**Table I.**
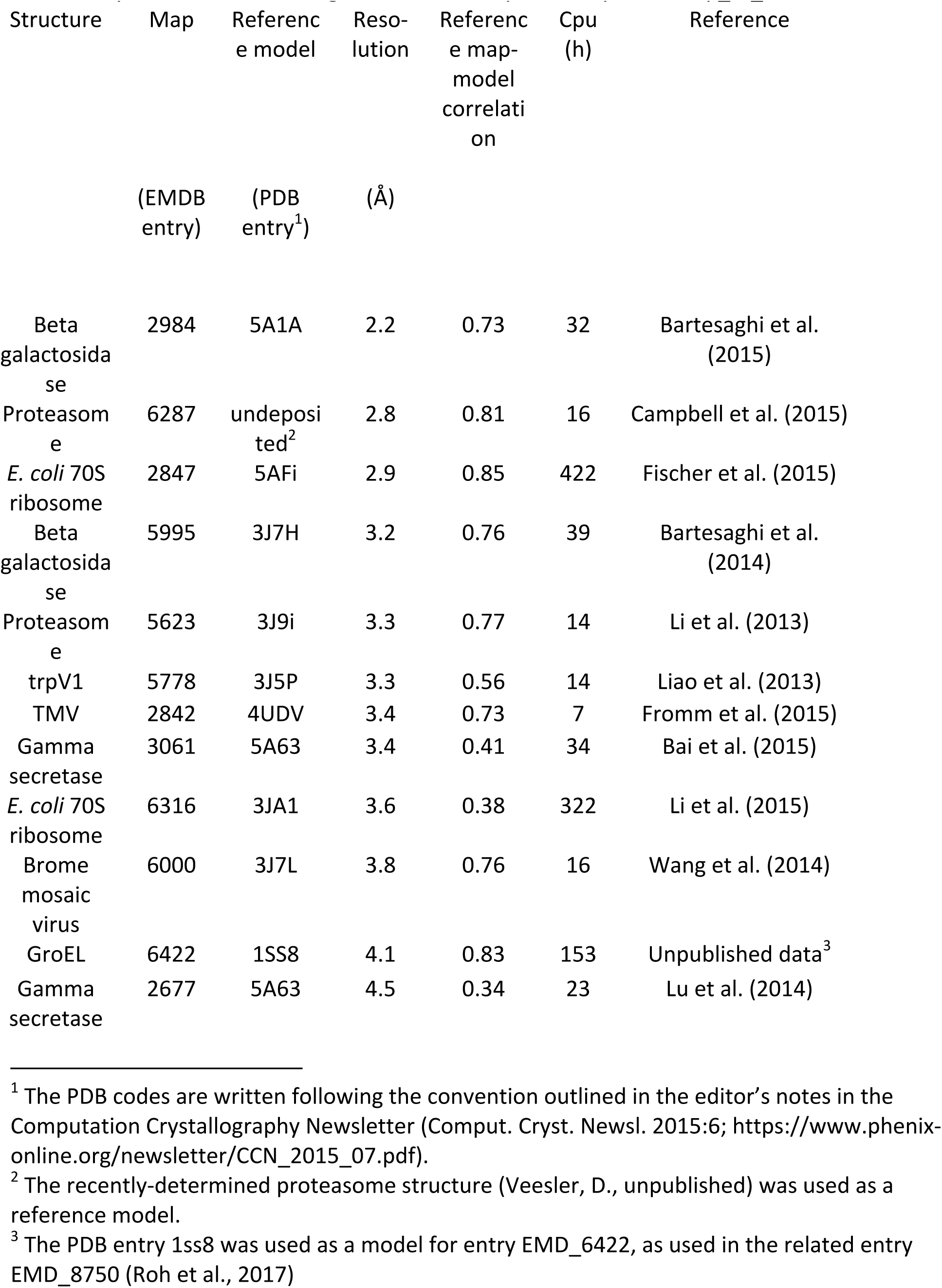
Cryo-EM Model Challenge structures analyzed with *phenix.map_to_model*

| Structure | Map | Reference model | Resolution | Reference map-model correlation | Cpu (h) | Reference |
| --- | --- | --- | --- | --- | --- | --- |
|  | (EMDB entry) | (PDB entry<sup>1</sup>) | (Å) |  |  |  |
| Beta galactosidase | 2984 | 5A1A | 2.2 | 0.73 | 32 | Bartesaghi et al. (2015) |
| Proteasome | 6287 | undeposited<sup>2</sup> | 2.8 | 0.81 | 16 | Campbell et al. (2015) |
| *E. coli* 70S ribosome | 2847 | 5AFi | 2.9 | 0.85 | 422 | Fischer et al. (2015) |
| Beta galactosidase | 5995 | 3J7H | 3.2 | 0.76 | 39 | Bartesaghi et al. (2014) |
| Proteasome | 5623 | 3J9i | 3.3 | 0.77 | 14 | Li et al. (2013) |
| trpV1 | 5778 | 3J5P | 3.3 | 0.56 | 14 | Liao et al. (2013) |
| TMV | 2842 | 4UDV | 3.4 | 0.73 | 7 | Fromm et al. (2015) |
| Gamma secretase | 3061 | 5A63 | 3.4 | 0.41 | 34 | Bai et al. (2015) |
| *E. coli* 70S ribosome | 6316 | 3JA1 | 3.6 | 0.38 | 322 | Li et al. (2015) |
| Brome mosaic virus | 6000 | 3J7L | 3.8 | 0.76 | 16 | Wang et al. (2014) |
| GroEL | 6422 | 1SS8 | 4.1 | 0.83 | 153 | Unpublished data<sup>3</sup> |
| Gamma secretase | 2677 | 5A63 | 4.5 | 0.34 | 23 | Lu et al. (2014) |

<sup>1</sup> The PDB codes are written following the convention outlined in the editor's notes in the Computation Crystallography Newsletter (Comput. Cryst. Newsl. 2015:6; https://www.phenix-online.org/newsletter/CCN_2015_07.pdf).
<sup>2</sup> The recently-determined proteasome structure (Veesler, D., unpublished) was used as a reference model.
<sup>3</sup> The PDB entry 1ss8 was used as a model for entry EMD\_6422, as used in the related entry EMD\_8750 (Roh et al., 2017)

### Results and Discussion

Fig. 1 illustrates the application of our map segmentation procedure (Terwilliger et al., 2018b) to the 2.9 Å cryo-EM reconstruction of the anthrax protective antigen pore (Jiang et al., 2015). The map has C7 symmetry (a 7-fold symmetry axis). Fig. 1A shows the 7-fold symmetry of the pore and illustrates one of the 7 chains in purple. Fig. 1B shows the density map with 7-fold symmetry. It can be seen that the density is much stronger for the extracellular region of the molecule than for the transmembrane part below. The 7-fold symmetry was used along with the map to identify symmetry-related regions of density in the map. Then a compact and connected unique set of density regions was chosen to represent the molecule. Fig. 1C shows the individual segmented regions of the map, and Fig. 1D shows the segmented region, augmented by neighboring regions of density.

**Figure 1.**
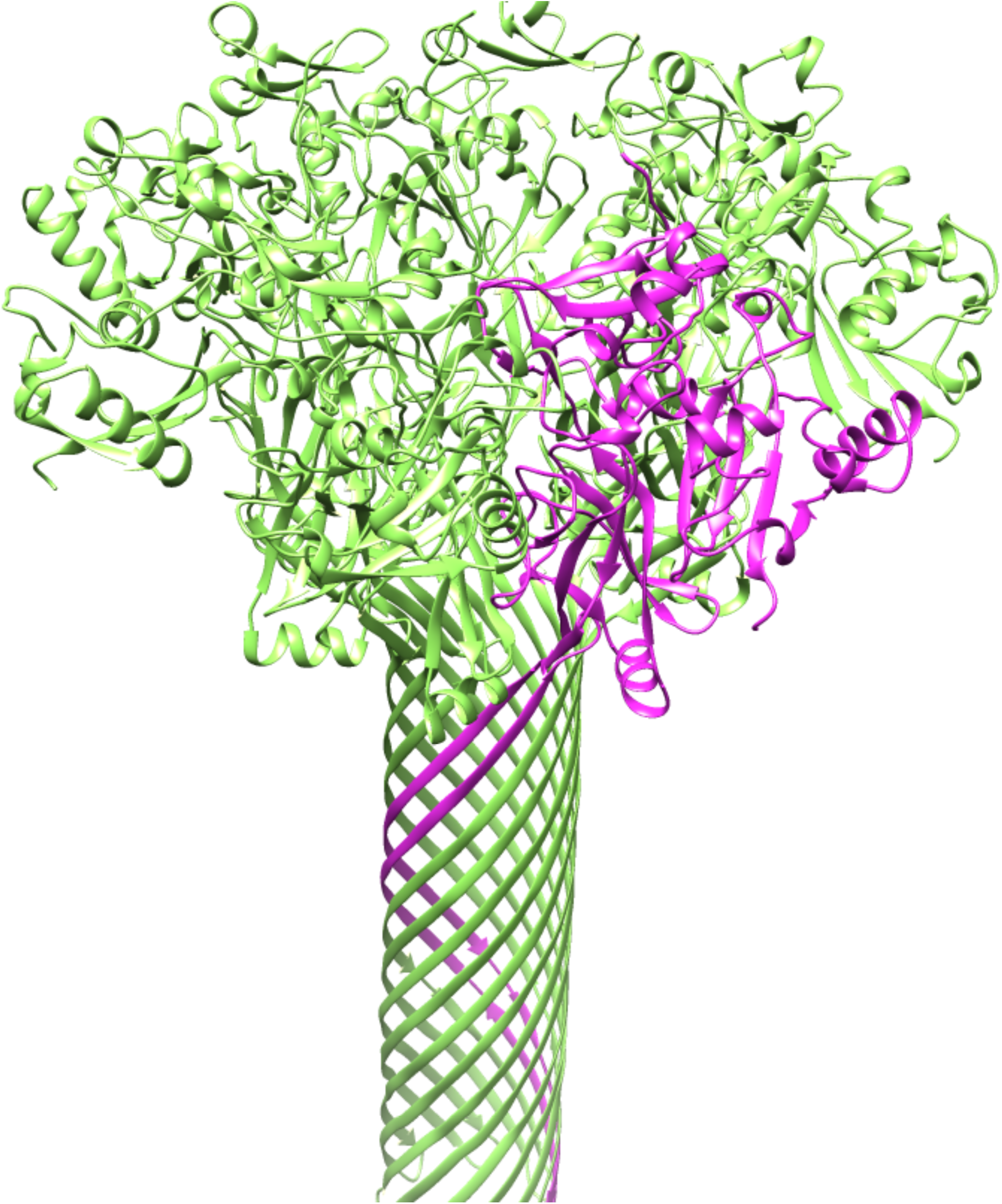
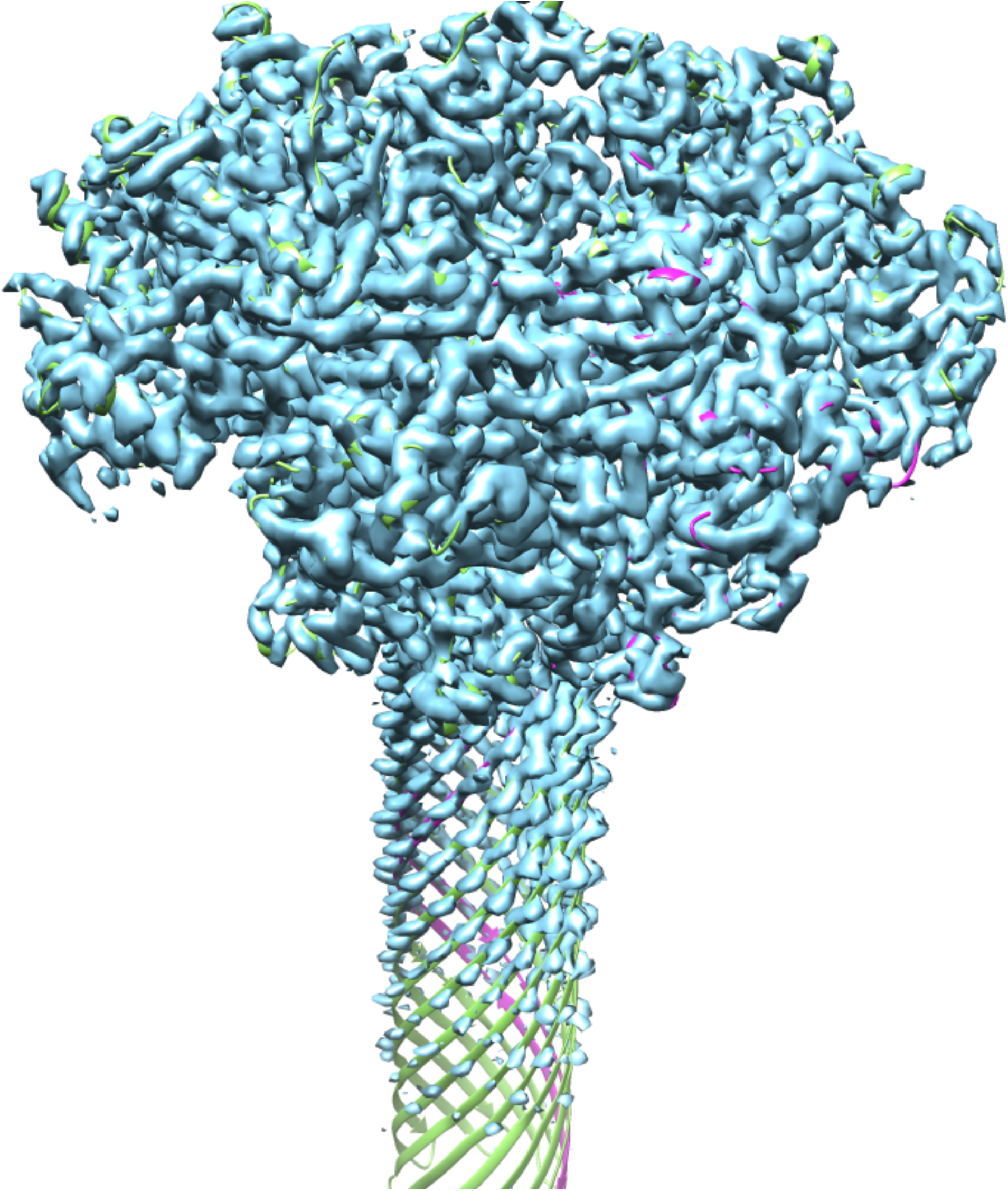
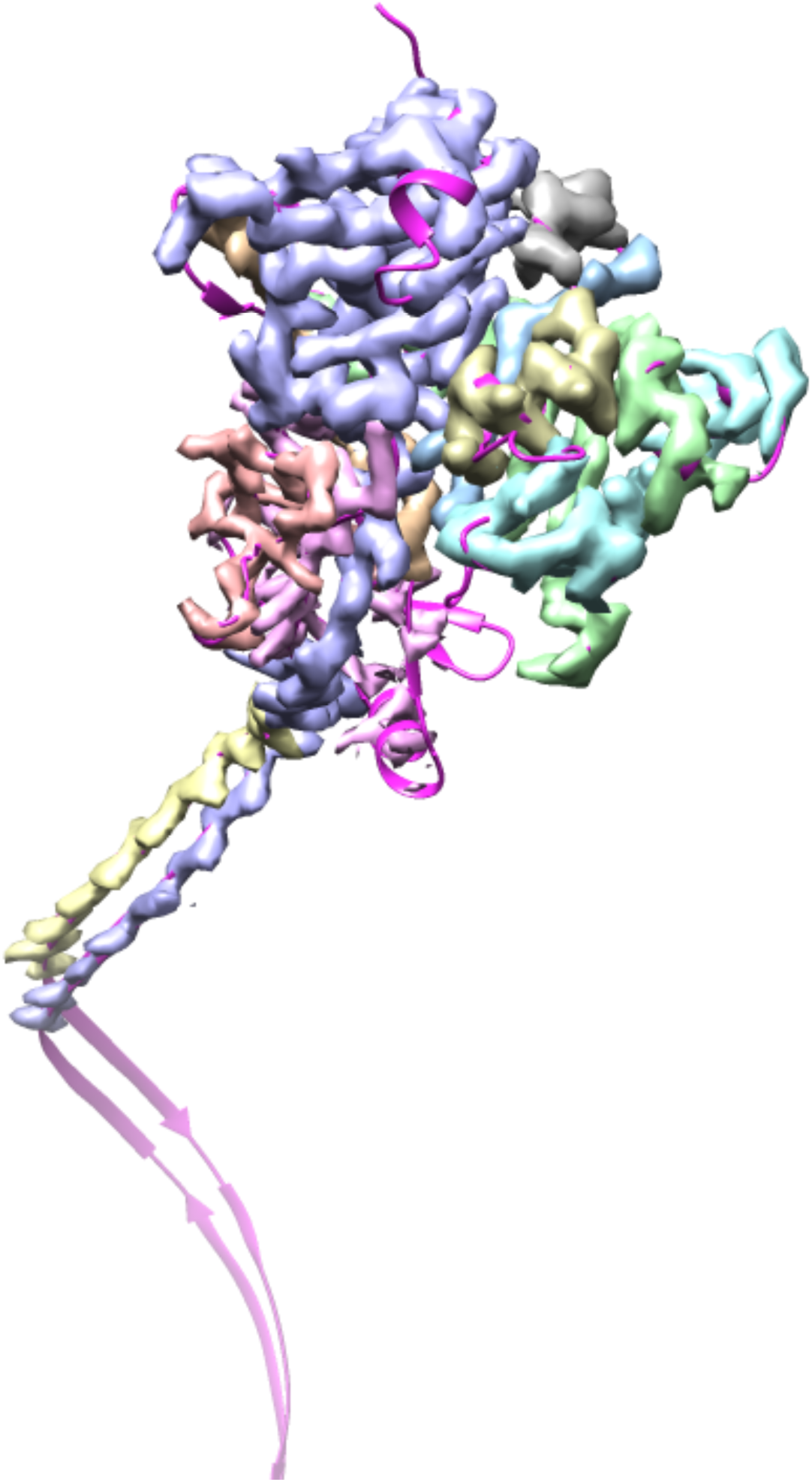
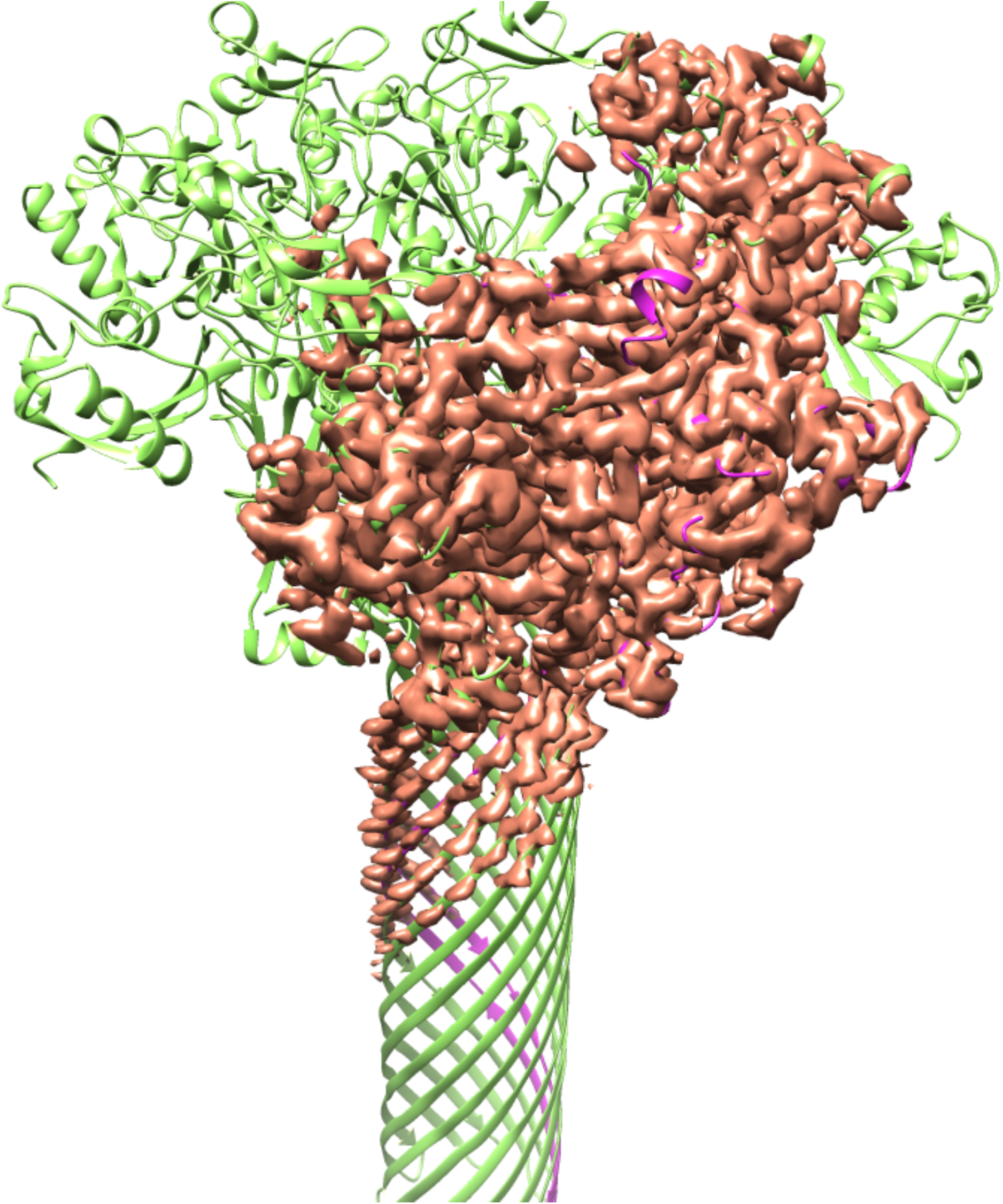
Segmentation of density for the anthrax protective antigen pore. A. Deposited structure of anthrax protective antigen pore with one of the 7 chains in purple. B. Density map illustrating the 7-fold symmetry used in the reconstruction. C. Individual segmented regions of the map superimposed on a single chain from the deposited structure. Note that the deposited structure was not used in the segmentation process. D. Illustration of the segmented region, augmented by neighboring regions of density.

We then applied our segmentation procedure to a large set of cryo-EM maps from the EMDB (Fig. 2). As expected, using the reconstruction symmetry of the maps in segmentation often resulted in a very large reduction in the volume that needed to be considered to include the unique part of each map (Fig. 2A). The average volume after segmentation and placing the unique segmented region in a new box was 8% of the starting volume of the maps. In most (206 of 218) of the cases illustrated in Fig. 2 we used the *add_neighbors* keyword to add a layer of regions around the unique molecular volume in order to increase the chance of finding a complete molecule. The 12 cases (EMD_2807, EMD_3137, EMD_5185, EMD_5600, EMD_6346, EMD_6630, EMD_6637, EMD_6688, EMD_8598, EMD_8605, EMD_8644, EMD_9518) where this was not done are those where the map was large (maps with 16M to 134M elements) and we attempted to keep the size of the region to be worked on to the minimum possible.

**Figure 2.**
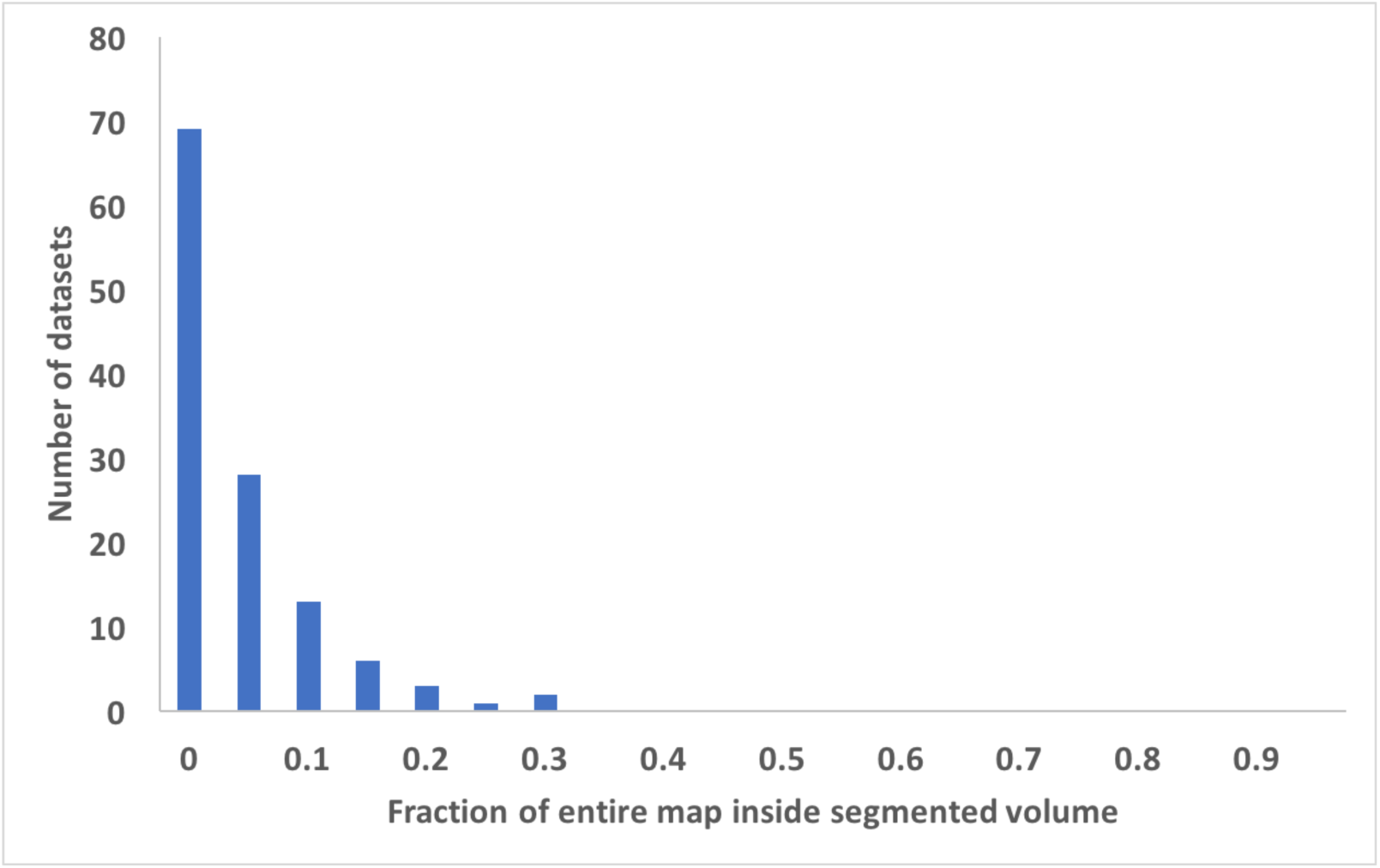
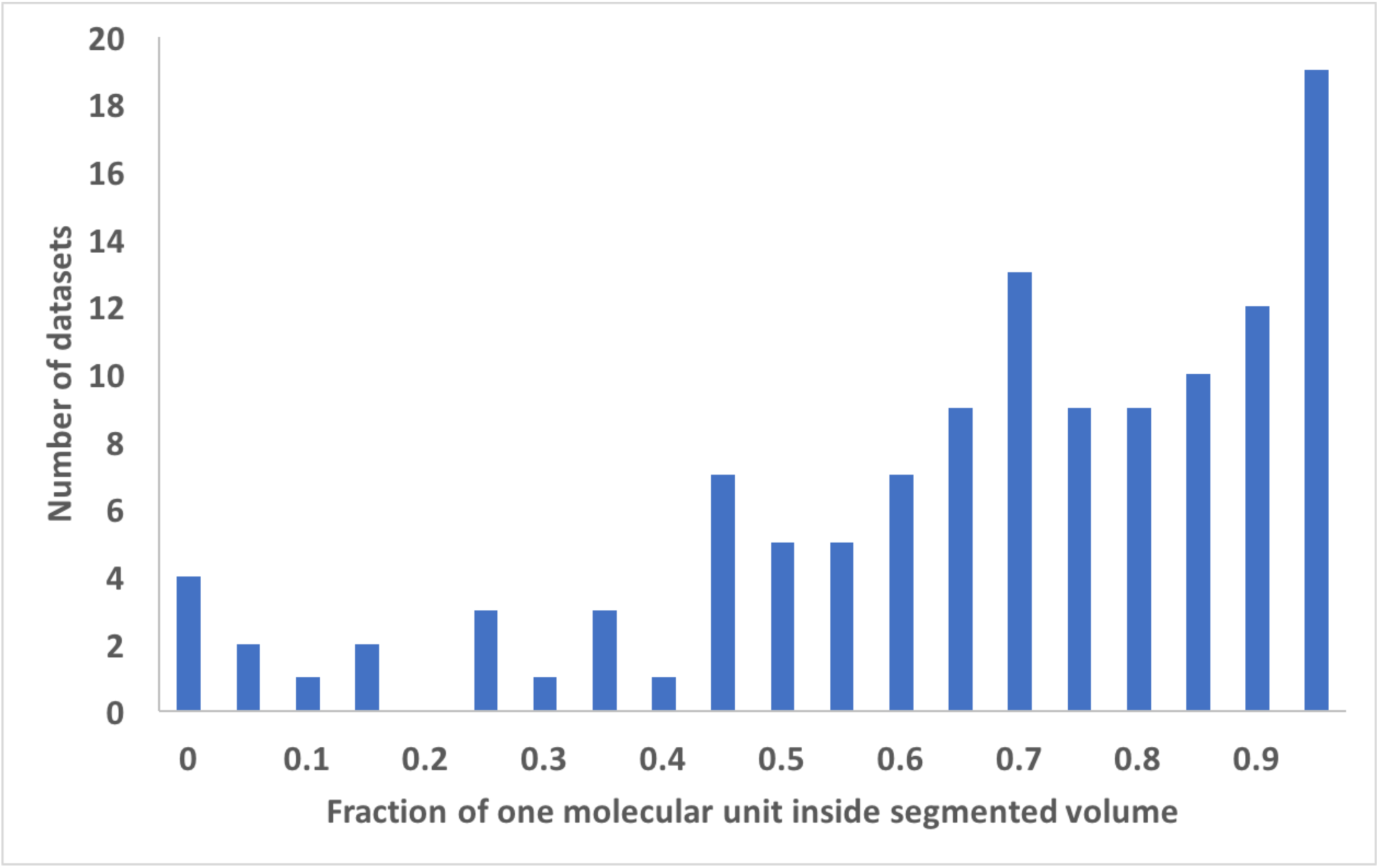
Histograms showing the results of application of the segmentation procedure to cryo-EM maps from the EMDB. Datasets are grouped according to (panel A), the fraction of original map required to represent the segmented region of each map, or (panel B), the fraction of each molecular unit contained within the segmented region of each map. In each panel, the label corresponds to the lower bound of each grouping. The values are grouped in increments of 0.05, so for example the number of datasets with values from 0.00 to 0.05 is shown over the ordinate of "0".

Fig. 2B illustrates the fraction of the unique molecular unit that is within the unique segmented region used for each map in Fig. 2A. This fraction of the molecule contained within the segmented region is estimated from the map-model correlation between the molecule and a map which is set to zero everywhere outside the segmented region, normalized to the map-model correlation without setting any of the map to zero. If the molecule is within the segmented region this normalized correlation will be unity, while if the molecule is split between different segmented regions it will be smaller. As shown in Fig 2B, the fraction within a single segmented region varies considerably among the 218 maps analyzed here, but the mean fraction was 0.72, indicating that typically a large fraction, but not all, of the molecular unit was contained within the segmented region.

We examined whether the fraction of the molecular unit contained within the segmented region (Fig. 2B) depended on the number of symmetry copies or the resolution of the map. The number of symmetry copies had only a small effect: maps with a single copy had an average fraction of 0.73 and maps with 60 copies had an average of 0.71. On the other hand, resolution had quite a substantial impact on the fraction within the segmented region: maps with resolution of 3.5 Å or better had a mean fraction of 0.82; maps with resolution of 4 Å or worse had a mean of 0.63.

We applied the combination of map sharpening, segmentation, and model-building as implemented in the *Phenix* tool *phenix.map_to_model* (Terwilliger et al., 2018b) to the 12 maps in the 2016 cryo-EM Challenge. The maps and corresponding reference models are listed in Table I along with the CPU hours required for the analysis, which ranged from 7 to 422 hours. Table II lists the number of residues that were built with C_α_ or P atoms within 3 Å of the corresponding atoms in the reference model by the *phenix.map_to_model* procedure, along with the fraction of the reference model represented by the model that was built and the fraction of residues that were assigned the correct residue identity. The number of residues built more than 3 Å from any residue in the reference model is also listed.

**Table II.** Results of Cryo-EM Challenge analysis with *phenix.map_to_model*

| Structure | Resolution |  | Residues in reference model | Built within 3 Å (Residues) | RMS distance (Å)<sup>1</sup> | Built (%) | Matched to sequence<sup>2</sup> (%) | Built further than 3 Å (Residues) |
| --- | --- | --- | --- | --- | --- | --- | --- | --- |
|  | (Å) |  |  |  |  |  |  |  |
| Beta galactosidase | 2.2 |  | 1022 | 842 | 0.6 | 82 | 75 | 10 |
| Proteasome | 2.8 |  | 422 | 327 | 0.6 | 78 | 59 | 3 |
| *E. coli* 70S ribosome | 2.9 | Protein | 6322 | 3212 | 0.7 | 51 | 23 | 2112 |
|  |  | RNA | 4748 | 2566 | 1.0 | 54 | 49 | 280 |
| Beta galactosidase | 3.2 |  | 1022 | 676 | 1.2 | 66 | 23 | 26 |
| Proteasome | 3.3 |  | 427 | 246 | 1.2 | 58 | 48 | 4 |
| trpV1 | 3.3 |  | 592 | 330 | 0.8 | 56 | 30 | 18 |
| TMV | 3.4 |  | 153 | 76 | 1.3 | 50 | 34 | 8 |
| Gamma secretase | 3.4 |  | 1223 | 832 | 1.0 | 68 | 24 | 80 |
| *E. coli* 70S ribosome | 3.6 | Protein | 7125 | 2479 | 2.0 | 35 | 8 | 3979 |
|  |  | RNA | 4685 | 1140 | 1.9 | 24 | 26 | 440 |
| Brome mosaic virus | 3.8 |  | 479 | 198 | 1.5 | 41 | 11 | 14 |
| GroEL | 4.1 |  | 524 | 309 | 1.3 | 59 | 16 | 10 |
| Gamma secretase | 4.5 |  | 1223 | 619 | 2.1 | 51 | 8 | 185 |

<sup>1</sup> The RMS distances for Cα/P atoms from the reference interpretations, only including the residues built within 3 Å of the reference model.
<sup>2</sup> The percentage of matched to sequence is the total number of residues in the automatically-built model correctly matched to sequence divided by the total number of residues in the reference model.

Overall, from 35% to 82% of the protein portions of the 12 structures were built within 3 Å of the corresponding reference models. For the two RNA structures, 24% and 54% of the RNA portions were built within 3 Å of the corresponding reference models. From 8% to 75% of the protein and RNA sequences were correctly assigned. For the non-ribosome structures, only a small proportion of the models built did not correspond at all to the deposited models. On the other hand, for the ribosome structures, a large fraction (over half for the 3.6 Å map) of the protein residues built did not correspond to the deposited models. Most of these incorrectly-built residues are located in regions that are RNA in the deposited models (recently we have developed a tool, *phenix.remove_poor_fragments* that can remove some of these incorrectly-built residues, but it was not available at the time of this work, TT, OS, PDA and PVA, unpublished). The models built by *phenix.map_to_model* have RMS distances for Cα/P atoms from reference interpretations of 0.6 Å to 2.1 Å.

The procedures developed here for map segmentation could be applied automatically to all of the 218 maps that we examined in the tests shown in Fig. 2. Further, all 12 of the maps in the 2016 Cryo-EM Model Challenge could be automatically sharpened, segmented and partially interpreted by the *phenix.map_to_model* procedure. It seems likely that combining the techniques developed here with other approaches for automatic model-building might lead to procedures that can automatically interpret an even larger part of cryo-EM maps.

## Acknowledgements

This work was supported by the NIH (grant GM063210 to PDA and TT) and the *Phenix* Industrial Consortium. This work was supported in part by the US Department of Energy under Contract No. DE-AC02-05CH11231 at Lawrence Berkeley National Laboratory. This research used resources provided by the Los Alamos National Laboratory Institutional Computing Program, which is supported by the U.S. Department of Energy National Nuclear Security Administration under Contract No. DE-AC52-06NA25396.Molecular graphics and analyses were performed with the UCSF *Chimera* package (Petterson et al., 2005) and with *Coot* (Emsley et al., 2010).

## References

Adams, P. D., Afonine, P.V., Bunkóczi, G., Chen, V.B., Davis, I.W., Echols, N., Headd, J.J., Hung, L.- W. Kapral, G.J., Grosse-Kunstleve, R.W., McCoy, A.J., Moriarty, N.W., Oeffner, R., Read, R.J., Richardson, D.C., Richardson, J.S., Terwilliger, T.C. & Zwart, P.H., 2010. PHENIX: a comprehensive Python-based system for macromolecular structure solution. Acta Cryst., D66, 213–221.

Afonine, P.V., Poon, B.K., Read, R.J., Sobolev, O.V., Terwilliger, T.C., Urzhumtsev, A., Adams, P.D. (2018a). Real-space refinement in *Phenix* for cryo-EM and crystallography. Acta Cryst D., in press.

Afonine, P.V., Klaholz, B.P., Moriarty, N.W., Poon, B.K., Sobolev, O.V., Terwilliger, T.C., Adams, P.D., Urzhumtsev, A. (2018b). New tools for the analysis and validation of Cryo-EM maps and atomic models. BioRxiv doi.org/10.1101/279844.

Baker, M.L., Yu, Z., Chiu, W., Bajaj, C., 2006. Automated segmentation of molecular subunits in electron cryomicroscopy density maps. J. Structural Biology 156, 432–441.

Baker, M.L., Ju, T., Chiu, W., 2007. Identification of secondary structure elements in intermediate-resolution density maps. Structure 15, 7–19.

Baker, M.L., Abeysinghe, S.S., Schuh, S., Coleman, R.A., Abrams, A., March, M.P., Hryc, C.F., Ruths, T., Chiu, W., Ju, T. (2011). Modeling protein structure at near atomic resolutions with Gorgon. J. Struct. Biol. 174, 360–373.

Bai, X.C., Yan, C.Y., Yang, G.H., Lu, P.L., Ma, D., Sun, L.F., Zhou, R., Scheres, S.H.W., Shi, Y.G. (2015). An atomic structure of human γ-secretase. Nature 525, 212–217.

Bartesaghi, A., Merk, A., Banerjee, S., Mathies, D., Wu, X, Milne, J.L., Subramaniam, S. (2015). 2.2 Å resolution cryo-EM structure of β-galactosidase in complex with a cell-permeant inhibitor. Science 348, 1147–1151.

Bartesaghi, A., Matthies, D., Banerjee, S., Merk, A., Subramaniam, S. Structure of β-galactosidase at 3.2-Å resolution obtained by cryo-electron microscopy. Proc. Natl. Acad. Sci. USA 111, 11709–11714.

Berman, H.M., Westbrook, J., Feng, Z., Gilliland, G., Bhat, T.N., Weissig, H., Shindyalov, I.N. and Bourne, P.E., 2000. The Protein Data Bank. Nucleic Acids Research, 28, 235–242.

Bernstein, F.C., Koetzle, T.F., Williams, G.J., Meyer, E.F., Jr., Brice, M.D., Rodgers, J.R., Kennard, O., Shimanouchi, T. and Tasumi, M., 1977. The Protein Data Bank: a computer-based archival file for macromolecular structures. J. Mol. Biol. 112, 535–542.

Campbell, M.G., Veesler, D., Cheng, A., Potter, C.S., Carragher, B. (2015). 2.8 Angstrom resolution reconstruction of the Thermoplasma acidophilum 20S proteasome using cryo-electron microscopy. eLife 4, e06380–e06380.

Cheng, A., Henderson R., Mastronarde, D., Ludtke, S., Schoenmakers, R.H.M., Short, J., Marabini, R., Dallakyan, S., Agard, D., Winn, M. (2015). J. Struct. Biol. 192, 146–150.

C. Colliex, C., Cowley, J. M., Dudarev, S. L., Fink, M., Gjønnes, J., Hilderbrandt, R., Howie, A., Lynch, D. F., Peng, L. M., Ren, G., Ross, A. W., Smith, V. H., Jr, Spence, J. C. H., Steeds, J. W., Wang, J., Whelan, M. J., Zvyagin, B. B. International Tables for Crystallography (2006). Vol. C, ch. 4.3, pp. 259–429.

Collins, P., Si, D. (2017). A graph based method for the prediction of backbone trace from cryo-EM density maps. ACM-BCB '17 Proceedings of the 8th ACM International Conference on Bioinformatics, Computational Biology, and Health Informatics.

Chen, M., Baldwin, P.R., Ludtke, S.J., Baker, M.L., 2016. De Novo modeling in cryo-EM density maps with Pathwalking. J. Structural Biol. 196, 289–298.

DiMaio, F., Chiu, W. (2016). Tools for model building and optimization into near-atomic resolution electron cryo-microscopy density maps. Methods. Enzymol. 679, 255–276.

Emsley, P., Lohkamp, B., Scott, W. G. & Cowtan, K. (2010). Features and development of Coot. Acta Cryst. D66, 486–501.

Fischer, N., Neumann, P., Konevega, A.L., Bock, L.V., Ficner, R., Rodnina, M.V., Stark, H. (2015). Structure of the E. coli ribosome-EF-Tu complex at <3 A resolution by Cs-corrected cryo-EM. Nature 520, 567–570.

Frenz, B., Walls, A.C., Egelman, E.H., Veesler, D., DiMaio, F., 2017. RosettaES: a sampling strategy enabling automated interpretation of difficult cryo-EM maps. Naure Methods 14, 797–803.

Fromm, S.A., Bharat, T.A., Jakobi, A.J., Hagen, W.J., Sachse, C. (2015). Seeing tobacco mosaic virus through direct electron detectors. J. Struct. Biol. 189, 87–97.

Jiang, W., Baker, M.L., Ludtke, S.J., Chiu, W., 2001. Bridging the information gap: computational tools for intermediate resolution structure interpretation. J. Mol. Biol. 308, 1033–1044.

Jiang J., Pentelute B.L., Collier R.J., Zhou, Z.H., 2015. Atomic structure of anthrax protective antigen pore elucidates toxin translocation. Nature 521, 545–549.

Kong, Y., Ma, J., 2003. A structural-informatics approach for mining beta-sheets: locating sheets in intermediate-resolution density maps. J. Mol. Biol. 332, 399–413.

Kong, Y., Zhang, X., Baker, T.S., Ma, J., 2004. A Structural-informatics approach for tracing beta-sheets: building pseudo-C(alpha) traces for beta-strands in intermediate-resolution density maps. J. Mol. Biol. 339, 117–130.

Lawson C.L., Patwardhan, A., Baker, M.L., Hryc, C., Garcia, E.S., Hudson, B.P., Lagerstedt, I., Ludtke, S.J., Pintilie, G., Sala, R., Westbrook, J.D., Berman, H.M., Kleywegt, G.J., Chiu, W. 2016. EMDataBank unified data resource for 3DEM. Nucleic Acids Res. 44, D396–D403.

Li, X., Mooney P, Zheng, S., Booth, C., Braunfeld, M.B., Gubbens, S., Agard, DA., Cheng, Y. (2013). Electron counting and beam-induced motion correction enable near-atomic-resolution single-particle cryoEM. Nat. Methods 10, 584–590.

Li, W., Liu, Z., Koripella, R.K., Langlois, R., Sanyal, S., Frank, J. (2015). Activation of GTP hydrolysis in mRNA-tRNA translocation by elongation factor G. Sci. Adv. 1, e1500169.

Liao, M., Cao, E., Julius, D., Cheng, Y. (2013). Structure of the TRPV1 ion channel determined by electron cryo-microscopy Nature 504, 107–112.

Lindert, S., Alenxander, N., Wotzel, N., Karakas, M., Stewart, P.L., Meiler, J., 2012. Em-Fold: De novo atomic-detail protein structure determination from medium-resolution density maps. Structure 20, 464–478.

Lu, P.L., Bai, X.C., Ma, D., Xie, T., Yan, C.Y., Sun, L.F., Yang, G.H., Zhao, Y.Y., Zhou, R., Scheres, S.H.W., Shi, Y.G. (2014). Three-dimensional structure of human gamma-secretase. Nature 512, 166–170.

Pettersen E.F., Goddard T.D., Huang C.C., Couch G.S., Greenblatt D.M., Meng E.C., Ferrin, T.E. 2004. UCSF Chimera-a visualization system for exploratory research and analysis. J Comput. Chem. 25, 1605–1612.

Pintilie, G. D., Zhang, J., Goddard, T. D., Chiu, W., & Gossard, D. C., 2010. Quantitative analysis of cryo-EM density map segmentation by watershed and scale-space filtering, and fitting of structures by alignment to regions. Journal of Structural Biology 170, 427–438.

Roh, S.H., Hryc, C.F., Jeong, H.H., Fei, X., Jakana, J., Lorimer, G.H., Chiu, W. (2017). Subunit conformational variation within individual GroEL oligomers resolved by Cryo-EM. Proc. Natl. Acad. Sci. U.S.A. 114, 8259–8264.

Terwilliger, T. C. (2003). Automated main-chain model-building by template-matching and iterative fragment extension. Acta Cryst. D59, 38–44.

Terwilliger, T. C. (2010a). Rapid model-building of ß-sheets in electron density maps. Acta Cryst. D66, 276–284.

Terwilliger, T. C. (2010b). Rapid model-building of α-helices in electron density maps. Acta Cryst. D66, 268–275.

Terwilliger, T. C. (2010c). Rapid chain-tracing of polypeptide backbones in electron density maps. Acta Cryst. D66, 285–294.

Terwilliger, T.C., Sobolev, O., Afonine, P.V., Adams, P.D. (2018a). Automated map sharpening by maximization of detail and connectivity. bioRxiv doi: https://doi.org/10.1101/247049.

Terwilliger, T.C., Adams, P.D., Afonine, P.V., Sobolev, O.V. (2018b). A fully automatic method yielding initial models from high-resolution electron cryo-microscopy maps. bioRxiv doi: https://doi.org/10.1101/267138

Volkmann, N. (2002). A novel three-dimensional variant of the watershed transform for segmentation of electron density maps. J. Struct. Biol. 138, 123–129.

Wang, Z., Hryc, C.F., Bammes, B., Afonine, P.V., Jakana, J., Chen, D.H., Liu, X, Baker, M.L., Kao, C., Ludtke, S.J., Schmid, M.F., Adams, P.D., Chiu W. (2014). An atomic model of brome mosaic virus using direct electron detection and real-space optimization. Nature Commun. 5 4808.

Wang, R. Y-R., Kudryashev, M., Li, X., Egelman, E.H., Basler, M., Cheng, Y., Baker, D., DiMaio, F., 2015. De novo protein structure determination from near-atomic-resolution cryo-em maps. Nature Methods 12, 335–341.

Yu, Z., Bajaj, C. (2008). Computational approaches for automatic structural analysis of large biomolecular complexes. IEEE/AC Transactions on computational bology and bioinformatics 5 568–582.

Zhang, Q., Bettadapura, R., Bajaj, C. (2012). Macromolecular structure modeling from 3D EM using VolRover 2.0. Biopolymers 97, 709–731.

Zhou N., Wang, H., Wang, J., 2017. EMBuilder: A Template Matching-based automatic model-building program for high-resolution cryo-electron microscopy maps. Scientific Rep. 7, 2664.

